# Coherent-hybrid STED: a tunable photo-physical pinhole for super-resolution imaging at high contrast

**DOI:** 10.1101/381343

**Authors:** António J. Pereira, Mafalda Sousa, Ana C. Almeida, Luísa T. Ferreira, Ana Rita Costa, Marco Novais-Cruz, Cristina Ferrás, Mónica Mendes Sousa, Paula Sampaio, Michael Belsley, Helder Maiato

## Abstract

Resolution in microscopy is not limited by diffraction as long as a nonlinear sample response is exploited. In a paradigmatic example, stimulated-emission depletion (STED) fluorescence microscopy fundamentally ‘breaks’ the diffraction limit by using a structured optical pattern to saturate depletion on a previously excited sample area. Two-dimensional (2D) STED, the canonical low-noise STED mode, structures the STED beam by using a vortex phase mask, achieving a significant lateral resolution improvement over confocal fluorescence microscopy. However, axial resolution and optical sectioning remain bound to diffraction. Here we use a tunable coherent-hybrid (CH) beam to improve optical sectioning, markedly reducing background fluorescence. CH-STED, which inherits the 2D-STED immunity to spherical aberration, diversifies the depletion strategy, allowing an optimal balance between two key metrics (lateral resolution and background suppression) to be found. CH-STED is used to perform high-contrast imaging of complex biological structures, such as the mitotic spindle and the neuron cell body.

## Introduction

Microscopy is conventionally limited to a resolution of approximately one half of the wavelength used to probe the sample. Some techniques have been developed which evade this optical ‘diffraction limit’ by harnessing the fact that fluorescence microscopy is not governed exclusively by optics laws, but instead involves the response of the fluorophore itself. In one particular case, stimulated emission depletion (STED) microscopy surpasses the diffraction limit by silencing the peripheral molecules of a previously excited spot, before they can spontaneously emit (Hell and Wichmann, 1994, Klar and Hell, 1999). Silencing is achieved by the process of stimulated emission and implemented by exposing the excited spot with a depletion beam (the ‘STED beam’) with a proper wavelength to induce stimulated emission and featuring a central dark spot surrounded by steep intensity gradients (Keller et al., 2007), thus forming a ‘doughnut’ shape. As a result, and if saturation is reached, only a subdiffraction sized spot of fluorophores will survive the action of the depletion beam and thus eventually fluoresce.

Usually assembled on a confocal microscope, the STED beam’s intensity, timing and polarization structure have to be engineered to form a sharp ‘intensity-zero’ at the focal point immediately after the sample has been excited (Galiani et al., 2012). Phase masks typically used to generate doughnuts are of two types. In the original design, the STED laser was modulated by a π-shifted ‘top-hat’ phase mask (z-STED mode) that elicits a strong axial confinement, which can still be further enhanced by using a 4Pi configuration (Dyba and Hell, 2002). Although very efficient in improving axial resolution, z-STED provides a lower lateral resolution than 2D-STED, while being relatively noisy due to the spurious signal emitted by secondary excitation lobes (‘ghosts’) that are not well covered by the z-STED beam geometry. In addition, an intensity-zero will only form if the central disc of the top-hat has the correct dimension relative to the microscope objective’s back-aperture (Keller et al., 2007, Heine et al., 2018) or, more precisely, relative to the effective aperture covered by the laser beam. A scale mismatch will unbalance the contribution of the π-shifted electric fields, leading to a filling of the central zero that depletes the signal of interest, compromising resolution and in particular the signal-to-noise ratio (SNR). Such calibration of the phase mask scale depends on imaging conditions that can be regarded as experimental variables, such as refractive index mismatch, observation depth and a switch of the objective (Heine et al., 2018), making z-STED particularly demanding for the end-user.

The two main limitations of z-STED (filling of the intensity-zero and ghost signals) persist as a noise contribution when the z-STED beam is incoherently added to a second beam to improve point-spread function (PSF) isotropy, defining what is sometimes called 3D-STED (Harke et al., 2008b). Whilst the ghost signals can still be partially depleted by the added second beam, the filling of the intensity-zero cannot be corrected by incoherent addition, which by definition adds individual intensities. Additionally, 3D-STED also poses severe challenges concerning the critical co-alignment of the two STED beams’ zeros at the nanometer scale.

More frequently, a ‘vortex’ phase mask (2D-STED mode) (Torok and Munro, 2004) is used because it provides maximal lateral (xy) resolution for a given STED laser power (Keller et al., 2007), in essence requiring only a circularly polarized beam with the proper handedness. A crucial feature of the vortex mask is that it does not possess a radial scale, thus avoiding the problem of scale mismatch. In fact, a general consequence of a scale-less mask is that in the presence of an arbitrary radial phase perturbation, such as spherical aberration or an axial misalignment in the microscope’s optics, a zero of intensity still forms in the beam’s center (**Supplementary Text**). Thus, for good reasons, the vortex is regarded as a standard in STED microscopy, particularly in SNR-demanding conditions, with its sharp and resilient zero facilitating the swift operation and high performance seen in STED microscopy today.

The feature that makes the zero in 2D-STED relatively immune to phase perturbations has the disadvantage that the vortex-generated zero is not constrained axially. Consequently, the beam does not deplete out-of-focus fluorescence, providing only confocal-like sectioning capacity. The effective excitation PSF is a thin needle along the optical axis. The profound consequences on the SNR can be visualized in a scenario where, by using an intense 2D-STED beam, the needle-shaped fluorescent source is made so thin that the number of molecules that remain undepleted is vanishingly small. This condition is eventually reached even if an ideal beam with a ‘perfect central zero’ is assumed. The usual recourse is to reduce STED laser power, which widens the parabolic dip, so that neighbor molecules are added to the fluorescent population. However, by reducing STED laser power, the needle-shaped source dilates in all planes, rescuing both focal and the unwanted background signal (in effect at the cost of lateral resolution). As a result, the usual protocol does not permit an SNR improvement other than that derived from the confocal pinhole selectivity.

Instead of reducing the STED beam power, here we describe a more effective approach that makes use of a modified, single-beam, doughnut that dilates the dip primarily in the focal plane, providing focus-specific signal selection. This can be viewed as a ‘photo-physical pinhole’ (Klar and Hell, 1999). Used together with the STED beam’s power, this focus-specific geometrical transformation of the depletion beam allows the microscopist to tune the acquisition’s SNR without fundamentally compromising lateral resolution.

## RESULTS

### Generation of a robust concave doughnut with a bi-vortex mask

To dilate the focal dip of the depletion beam, a phase plate is first defined containing a vortex that is slightly contracted (*radius*= *ρ*) relative to the typical full-NA vortex (*radius*=*1*), which generates a correspondingly wider doughnut, as dictated by diffraction (**Fig. 1a**). Such dilation is global and must be somehow counteracted to achieve focus-specificity. With this aim, the peripheral area exposed by vortex contraction (a thin ring) is filled with a second vortex, with the same handedness, but out-of-phase. A regular thin ring aperture is known to generate an elongated and narrow spot (Born and Wolf, 2013, Sheppard and Hegedus, 1988). A thin vortex ring would then, by its own, generate an elongated and narrow dip (Gong et al., 2010), reminiscent of an inverted Bessel beam (Wang et al., 2017). When used together (coherent combination), the compound ‘bi-vortex’ mask generates a long narrow dip, slightly dilated specifically at the focal plane region (**Fig. 1a**). This dilated dark focal spot can be seen as arising from the destructive interference of the beam crests that would be generated by the individual vortices, providing the key ingredient of the presented method. The volume of the dip can be modulated by changing the bi-vortex *radius* ρ, smoothly approaching the 2D-STED geometry when ρ tends to 1 (or 0).

**Fig. 1.**
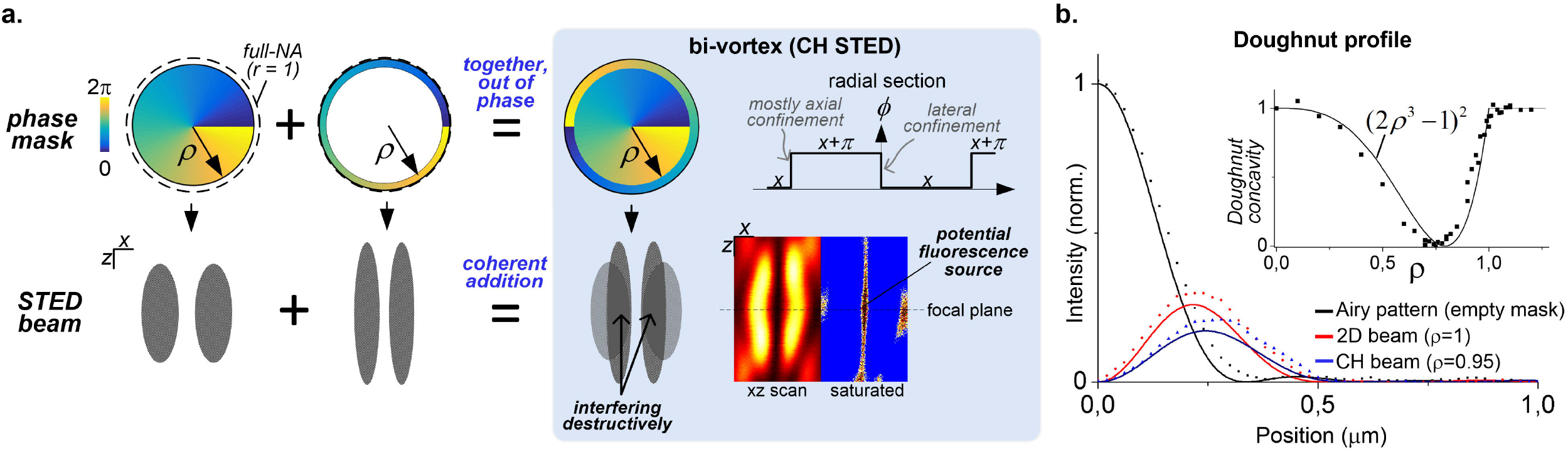
CH-STED beam: generation and focal profile. (**a**) Decomposition of the bi-vortex phase mask into two vortex components (inner disc and outer ring) provides a pictorial interpretation of the focus dip generation in CH-STED. A scattering image of a *xz*-scanned gold bead is shown using both a linear and a saturated look-up table (LUT), the latter providing a heuristic ‘preview’ of the undepleted region (‘potential fluorescence source’) geometry at high saturation. A bi-vortex radial section is shown to highlight the centered phase step (2D-STED) along with the additional off-axis steps, which provide the axial confinement component of CH-STED. (**b**) Experimental data-points and paraxial theoretical curves (solid lines) for the depletion beam focal profile. The single adjusted parameter of theoretical curves is the vertical scale, using the Airy pattern for normalization. In the inset, the bi-vortex radius is varied to show evolution of the beam’s geometrical confinement (concavity) along with the theoretical prediction. Experiment-theory comparison is not dependent on fitting parameters.

We call the beam created by the bi-vortex mask a coherent-hybrid (CH-)STED beam, as it can be viewed as the addition of a 2D-STED mask to a rescaled z-STED mask. Crucially, the phase of the bivortex mask (*ϕ*) may be viewed as a superposition of a radial function (a step function) with a vortex. This ‘radial vortex’ condition, *ϕ* = *f* (*r*) + *θ*, where *θ* is the azimuthal angle, is a sufficient condition for immunity of the zero to spherical aberration, mask scale or refractive index mismatches (**Supplementary Text**). In summary, radial aberrations, as well as instrumental imprecisions in setting *ρ* do not compromise the zero, making the CH-STED beam operationally rugged.

### CH-STED beam profile and the fluorescent spot after depletion

To appreciate the effect of the bi-vortex on lateral resolution, an analytical expression for the depletion beam intensity profile was derived under the paraxial condition (**Supplementary Text**). Although vectorial diffraction is required for a rigorous analysis (Mansuripur, 1986), assuming sufficient symmetry conditions (as imparted by the use of circular polarization) allows this approximation to deliver a quantitative insight into the effect of *ρ* on resolution. The intensity profile at the focal plane (**Fig. 1b**) in the neighborhood of the optical axis was found to follow *I* ∝(1 − 2*ρ*^3^)^2^ *x*^2^ (**Fig. 1b**, inset), where *x* is the distance to the optical axis.

We experimentally tested whether the sought-after axial performance of CH-STED would compensate for the effect of *ρ* on lateral resolution. Using a confocal-based pulsed-STED system at a 775nm depletion wavelength, 40nm-diameter fluorescently-labeled nano-beads (Crimson beads, excitation 640nm) were scanned in *xz*. The CH-STED PSF was observed to follow the behavior anticipated by inspection of the STED beam shape, namely it provides a degree of axial constrain (absent in 2D-STED) and a very efficient depletion of the ghost spots as compared to z-STED (**Fig. 2a**). Essentially, reshaping of the STED beam changed the curvature sign (from convex to concave) of the contour lines around the node, revealing the qualitative shift of the CH-STED when compared to the mere change of power in 2D-STED (**Fig. 2b**). The effect of the CH-STED beam on fluorescence is thus a suppression of out-of-focus signal, leaving a very low-noise ampoule-shaped PSF (**Fig. 2b**, bottom), which should be more suitable than 2D-STED for imaging thick and complex environments.

**Fig. 2.**
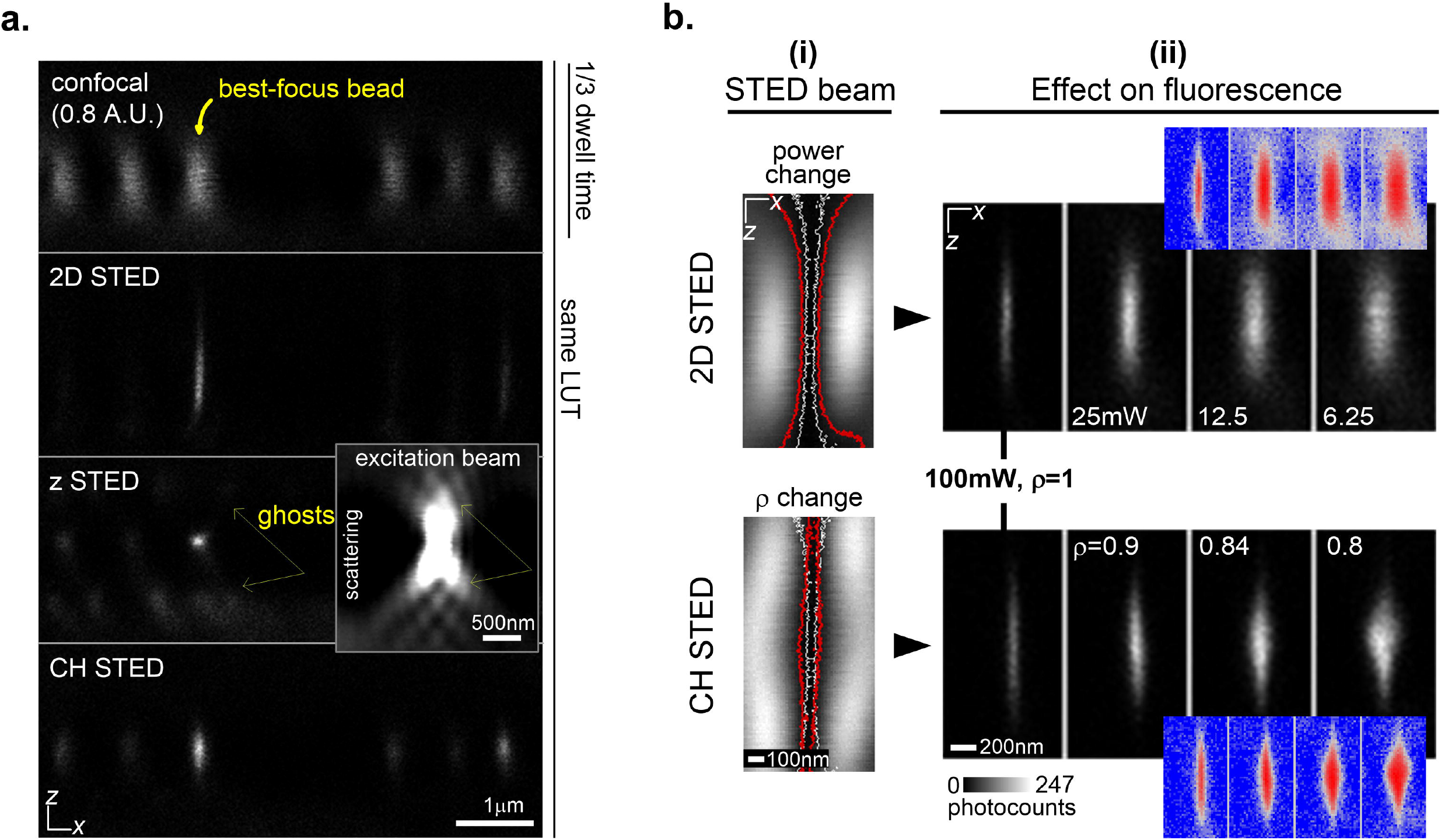
STED-modes PSFs. (**a**) Nano-bead fluorescence xz-scans in different STED modes (same LUT display). The inset highlights the origin of the excited ghosts that are poorly depleted by the z-STED beam. (**b**) (i) Gold bead scattering cross-section in 2D-STED (top) and CH-STED (bottom) with isophote lines defining the saturation contours. White-to-red transition represents signal rescue behavior in 2D-STED (STED power change, top) and in CH-STED (ρ change, bottom). (ii) Effective PSF in the two STED regimes displayed in (i). The four steps shown in each path were chosen so that the lateral in-focus resolution levels display a similar evolution in the top and bottom rows. Red-blue images (same LUT) are log-transformed versions to highlight noise suppression performance.

### Cell imaging upon switching to the bi-vortex CH-STED

To test the performance of CH-STED, we chose a complex biological object: the mitotic spindle. In the mitotic spindle, we observe that signal from interpolar microtubules and kinetochore fibers (a microtubule bundle that is attached to the kinetochore region of chromosomes), which are immersed in the noisy environment of the spindle, emerges after entering the CH-STED mode (**Fig. 3a**). Astral microtubules, as well as interphase microtubules, which are more isolated and do not form bundles, show the expected loss in lateral resolution, but even here microtubules generally become more visible due to the decrease in the defocused signal emitted from neighboring planes, providing better contrast (**Fig. 3a,b, Supplementary Fig. S1**). Importantly, this occurs without any change in the other acquisition settings, such as STED beam power. Indeed, additional STED power could be provided to increase lateral resolution upon switching to CH-STED, but this is not a general requirement when imaging thick or noisy environments.

**Fig. 3.**
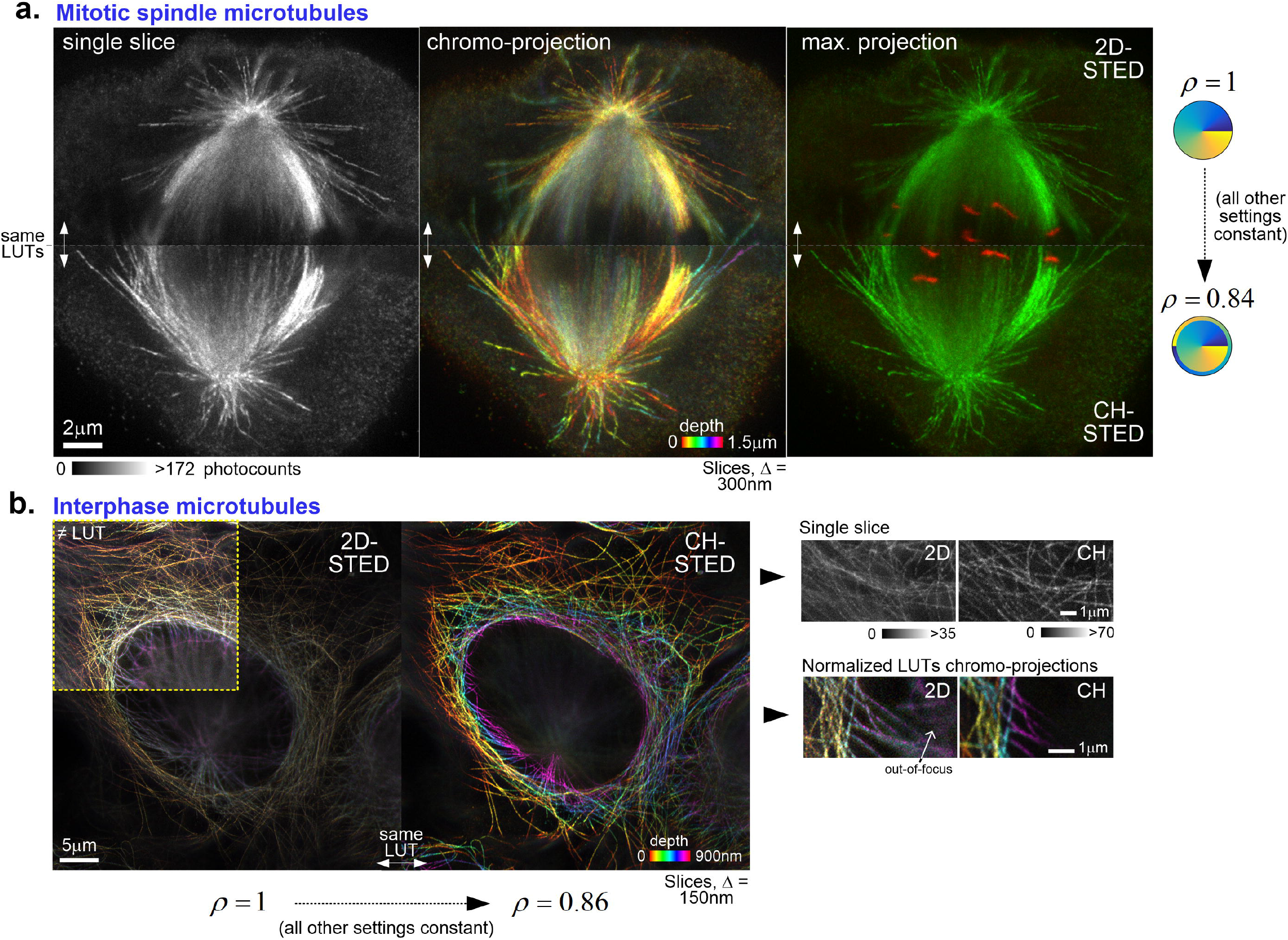
Direct comparison between 2D-STED and CH-STED by morphing the STED beam only. (**a**) Single-slice and projections obtained from a z-stack acquisition of a tubulin-labeled Indian muntjac mitotic spindle, along with a kinetochore marker (CREST) in the last panel. Top- and bottom-halves were acquired at vertically adjacent ROIs with all acquisition parameters kept constant except the bivortex radius, *ρ*, which was changed from >1 (2D-STED) to 0.84 (CH-STED). Pixel size, 30nm. Slice distance, 300nm. STED laser power, 80mW. (**b**) Z-stack acquisition chromo-projection of a tubulin-labeled U2OS interphase cell. Small ROIs are shown (right) as single slice and chromo-projections to highlight STED performance in areas with overlaid/tilted microtubule arrays. Pixel size, 35nm. Slice distance, 150nm. STED laser power, 80mW.

### Characterization of the ampoule-shaped CH-STED PSF

To more objectively assess CH-STED performance, we used basic metrics for analyzing the microscope’s PSF in the optical sectioning range, *ρ* ≈ 0.8−1 (**Fig.4a, see also Fig. 1b**) and at a varying STED power level. To establish metrics for the sought-after axial confinement, PSF width was measured at the focal plane (D_0_, **Fig. 4b**) and at a plane defocused by one Rayleigh range (*z_R_*=260nm), a standard measure of the half-depth of focus for a Gaussian beam (**Fig. 4b**). The PSF undergoes the usual scale transformation at varying STED power (**Fig. 4c**). For CH-STED, a shape transformation is observed towards increased confinement when *ρ* is decreased, as suggested by the intersection of the D_0_ and D_z_ curves (**Fig. 4d**). The lateral resolution data is in agreement with theoretical calculations (solid line in **Fig. 4d, Supplementary Text**), where a parabolic approximation to the depletion beam proves insufficient. This reflects the fact that, in CH-STED, power and resolution are not univocally related, lending the fluorescent profile sensitive to the detailed structure of the depletion beam whenever high STED power is used for high sectioning (i.e., lower lateral resolution, *ρ* < 0.9) (**Supplementary Text**).

**Fig. 4.**
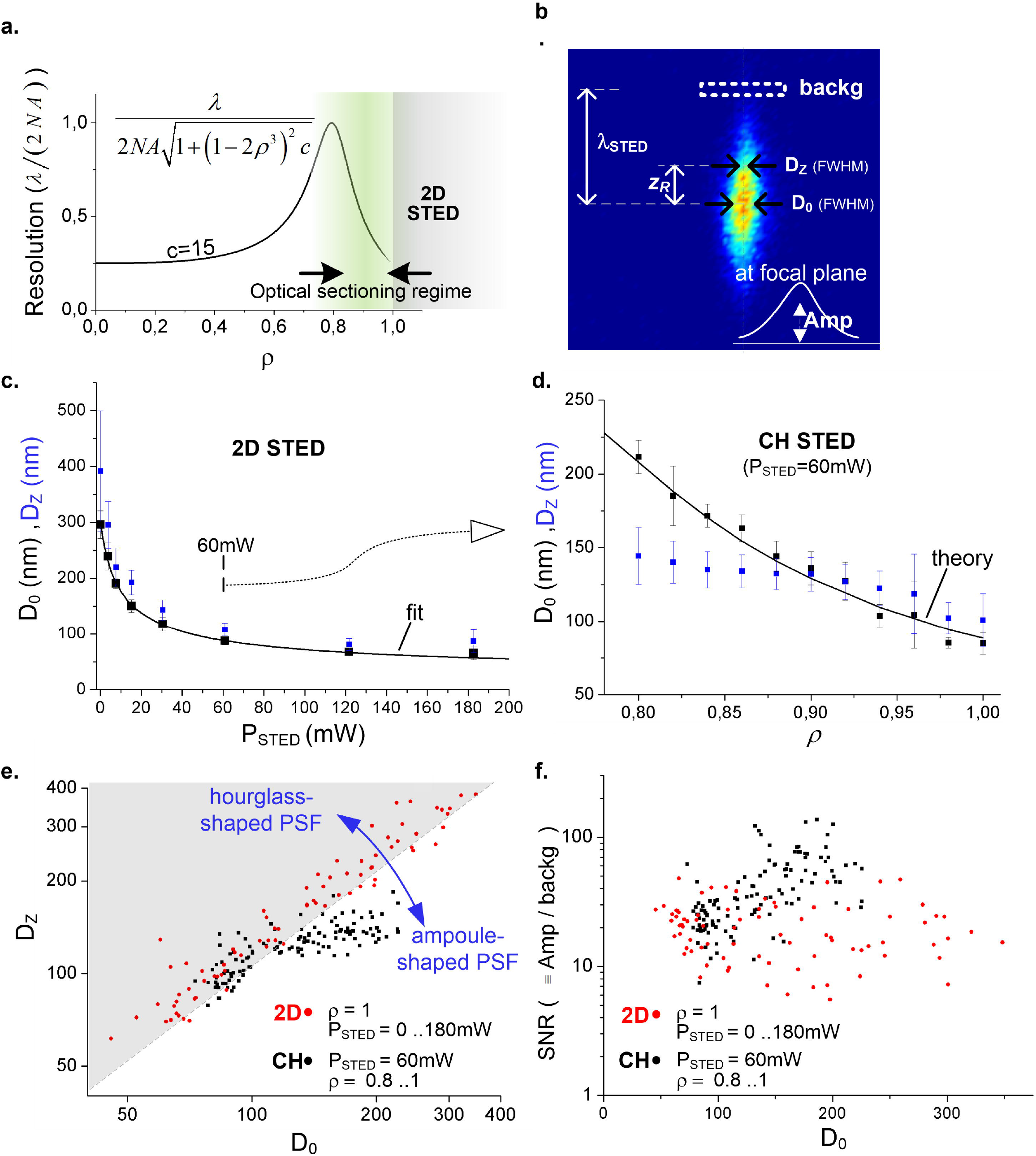
STED modes morphology and background suppression using 40-nm fluorescent beads. (**a**) Bivortex theoretical resolution (in the parabolic approximation) at a saturation value (*c*=15) where 2D STED displays a four-fold resolution improvement. (**b**) Metrics for assessing the performance of the STED modes, where the defocused plane chosen for measuring confinement is at a Rayleigh range distance (*z_R_*=260nm) from the focal plane. The large-defocus plane used for background estimation is at a *λ_STED_* distance (≈780nm=3*z_R_*) from the focal plane. (**c**) PSF lateral dimension (mean±s.d., n=10 beads per datapoint) at and away from the focal plane in 2D-STED as a function of STED laser average power. (**d**) PSF lateral dimension (mean±s.d., n=10 beads per datapoint) at and away from the focal plane in CH-STED as a function of *ρ*, using an intermediate-range STED power (60mW). (**e**) Scatter plot comparison of 2D-STED and CH-STED (at 60mW) using a common parameter (*D_0_*) as the independent variable. Each datapoint represents one bead. Axes were cropped at 400nm, leaving three 2D-STED (red) datapoints not displayed (used and accounted for in quantifications in **Fig. 4c**). (**f**) Background suppression estimation using the in-focus Gaussian curve fit amplitude (defined in (b)) relative to an average value at the background ROI (defined in (b)).

To enable a more direct comparison between 2D-STED and CH-STED performances, *D_Z_* was measured as a function of a common independent variable - lateral resolution, *D_0_*. In these scatter plots (**Fig. 4e**), the instrumental parameters *P_STED_* and *ρ* are implicit variables that probe the *D*_0_ × *D_z_* space. Here, the bottom-right half-space is populated by experimental PSFs that get narrower away from the focus (‘ampoule-shaped’), as opposed to the hourglass-shaped PSF typical of confocal and 2D-STED microscopes. In addition to the progressive geometrical confinement of the CH-STED PSF, a strong attenuation is observed for the integrated signal emanated from off-focus planes (**Fig. 4f**), as measured by the PSF amplitude relative to a background signal measured at a λSTED axial distance (=3_Z_R__) (defined in **Fig. 4b**).

### Exploration of the three CH-STED basic operation modes in cells

Thus far, we used either *P_STED_* or *ρ* modulation to tune the PSF (**Fig. 2b, Figs. 3a,b**). However, as independent and instrument-set parameters, the full two-dimensional space they define can be easily explored. Using the simplified parabolic dip approximation (**Fig. 2b**) and a first-order approximation to the depletion process (Harke et al., 2008a), the combined effect of ρ and *P_STED_* on lateral resolution is approximately given by

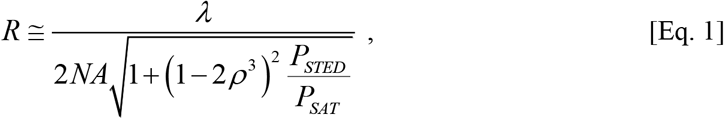

where *P_SAT,_* a saturation power, sets the scale for resolution improvement. Eq. 1 should be used as a rough resolution scale for ρ-values higher than ≈0.85 (**Supplementary Text**). As expected, a third (‘constant-resolution’) mode arises naturally (**Fig. 5a**) through the combination of a decreasing ρ with an increased power (Eq. 1). Here, the extra power not only recovers lateral resolution but now provides an active optical sectioning improvement by narrowing down the nodal line along the optical axis, which amounts to an improved axial resolution.

**Fig. 5.**
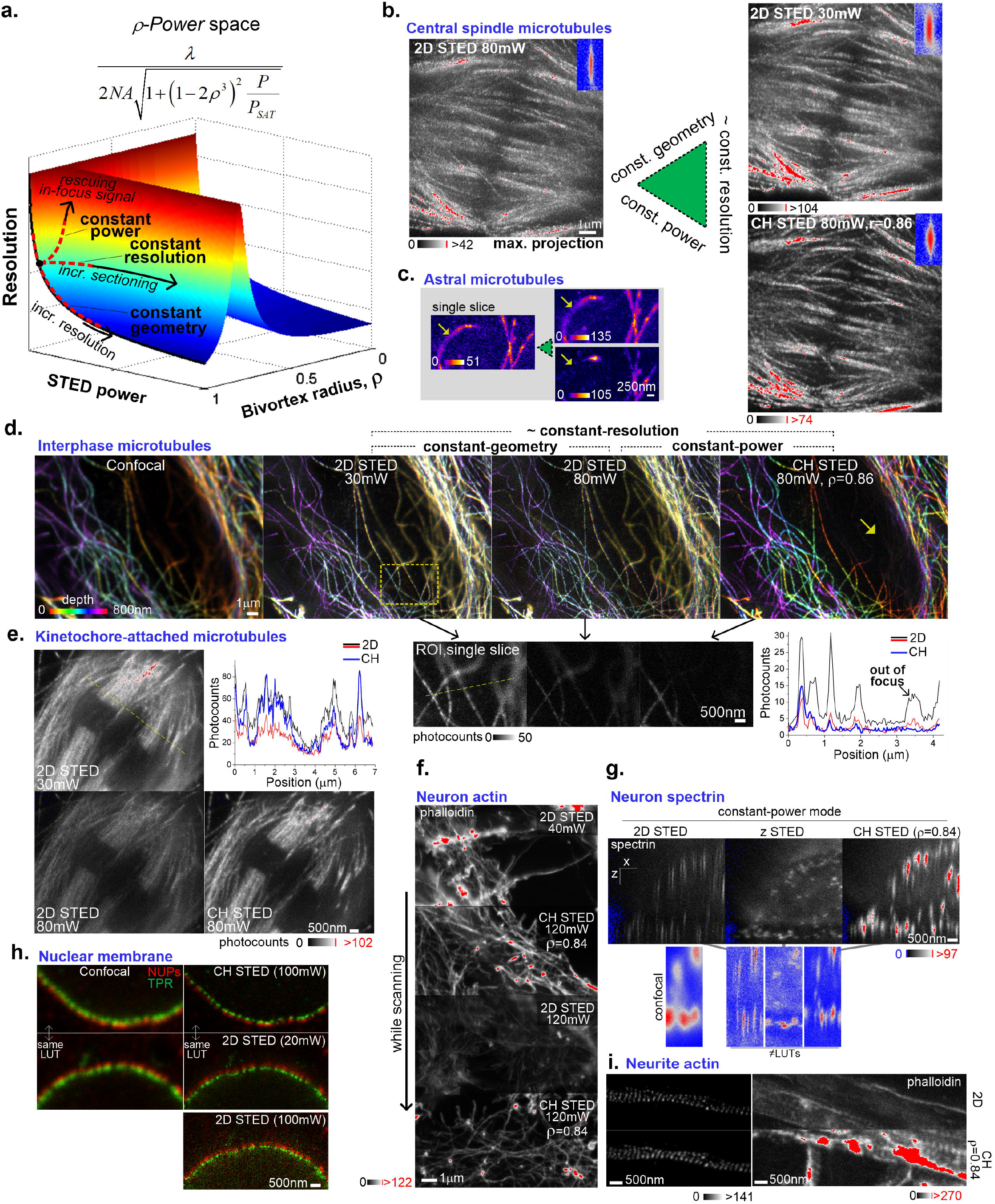
2D-STED vs. CH-STED in pairwise combinations. (**a**) Surface displaying theoretical resolution (under a parabolic approximation) across the *ρ -Power* parametric space. The three dimensions provide three natural modes for instrument operation: constant-power, constant-geometry (of which 2D-STED is one particular case) and constant-resolution. (**b**) Maximum-intensity projection of an anaphase central spindle in a U2OS tubulin-labeled cell. Blue-red insets (top-right corner) are pictorial examples (taken from **Fig. 2b**) of the PSF cross-section qualitative differences (not related to the actual acquisition). (**c**) Single slice of astral microtubules in a prometaphase U2OS cell showing suppression of defocused portions of a tilted microtubule when imaged in CH-STED. (**d**) Chromo-projection from a z-stack acquisition of an interphase Indian Muntjac cell. Color definition is a readout for optical sectioning, which can more objectively be assessed on the single slice close-up and associated intensity profile (taken from the yellow dashed line; black and red profiles are for the 30mW and 80mW 2D-STED images), where defocused signal is seen to be more efficiently suppressed than in the other STED modes. Pixel size, 30nm. Slice distance, 200nm. CH STED *ρ* =0.86. (**e**) Central region of tubulin-labeled Indian Muntjac mitotic spindle showing kinetochore-microtubule fibers attached to sister kinetochores. Intensity profile lines are shown for the yellow dashed line (black and red profiles are for the 30mW and 80mW 2D-STED images), Pixel size, 20nm. (**f**) Live shift across three STED modes during scanning of a neuron soma region, with phalloidin-labeled actin. Pixel size, 25nm. (**g**) 2D-, z- and CH-STED comparison at constant power by xz-scan of a neuron soma labeled with βII-spectrin. A small region is shown with auto-scaled LUTs along with the confocal acquisition. (**h**) Dualcolor STED (excitations at 560 and 640nm) of nuclear pore components in HeLa cells. Pixel size, 35nm. (**i**) Constant-power acquisitions on an actin-labeled isolated axon (left) and on less-isolated neurites (right).

To standardize CH-STED and 2D-STED comparative imaging using biological specimens, a three-acquisition sequence was followed, allowing a pair-wise comparison of the defined base-modes: constant-geometry, constant-power and constant-resolution (**Fig. 5a**). Different biological contexts (**Fig. 5b-h**) indicate that entering the CH-STED regime provides background rejection at high focal signal level. This is shown by the fact that intensity profiles (line-plots in **Figs. 5d,e**) display a high fluorescence level relative to the constant-power counterpart (the high-power 2D-STED) and a low background relative to the constant-resolution counterpart (the low-power 2D-STED), which amounts to a stretched dynamic range. As opposed to isolated structures, noisier environments, such as perinuclear areas of interphase cells (**Fig. 5d** and **Fig. 3b**)), mitotic cells (**Fig. 5b,e** and **Fig. 3a**), tilted microtubules (**Fig. 5c** and **Fig. 3b**), the actin or βII-spectrin distribution in the neuron soma (**Fig. 5 f,g**), nucleoporins (**Fig. 5h**) or even less isolated neurites (**Fig. 5i**, right), deliver increased structural information when observed in the CH-STED regime. The exception is very thin or isolated objects, such as peripheral interphase microtubules or isolated axons lying on the coverslip (**Fig.4i**, left), where optical sectioning is provided by the sample itself.

## DISCUSSION

Resolution improvement demands smaller localized excitation volumes and, consequently, faces a decreased molecule population per pixel. Thus, the pursuit for super-resolution imaging is often accompanied by a battle for signal over noise. In the case of STED microscopy this poses specific implementation challenges (Bianchini et al., 2015), pushing continuous developments (Vicidomini et al., 2018) such as pulse synchronization and gated-detection (Leutenegger et al., 2010, Vicidomini et al., 2013), two-photon excitation (Moneron and Hell, 2009, Takasaki et al., 2013) or beam polarization engineering (Reuss et al., 2010, Xue et al., 2012). Here, deviating from a search for best resolution (Keller et al., 2007), it is suggested that a metric combining resolution and background noise suppression be considered, a heuristic that seems better adapted to the ampoule-shaped CH-STED PSF than the typical hourglass-shaped PSF (**Fig. 2b**). Working in series with confocal pinhole (passive) sectioning, ampoule-shaped beams (or in general, optical bottles (Arlt and Padgett, 2000)) behave as ‘photo-physical pinholes’ (Klar and Hell, 1999) that actively suppress out-of-focus signal at its origin.

z-STED is one example of a photo-physical pinhole that provides unsurpassed axial resolution. However, it displays relatively low lateral confinement and is highly susceptible to imperfections in generating the zero. It is thus reasonable to search for a quasi-2D STED beam, in the sense of inheriting its high-resolution and high-quality zero, but behaving also as a pinhole. The CH-STED beam accomplishes this as soon as *ρ* is decreased below 1. Unless sub-diffraction sectioning is provided by the sample itself (Punge et al., 2008) or by the use of evanescent waves, such as in TIRF-STED (Gould et al., 2011), it seems unlikely that the exact value *ρ*=1, which defines 2D-STED, is the best choice in any given context within imaging-based, time-domain (e.g. FCS, Eggeling et al., 2013) or litography (Wollhofen et al., 2013) STED variants.

CH-STED does not require modifications in the (circular) polarization state of the STED beam used in typical setups, and thus can be implemented in SLM-based systems, which readily provide the required flexibility. The often-used static phase plates (e.g. polymer-based) could however perform adequately by using a ‘zoom’ optical system that persistently images a bi-vortex plate at the back focal plane of the objective at a variable magnification. Independent of implementation details, a CH-STED beam can still be incoherently combined to a z-STED beam to provide a more isotropic resolution, despite the usual technical demands in terms of alignment and aberrations control (Patton et al., 2015, Heine et al., 2018).

Still to be explored is the question whether in CH-STED (or its incoherent combinations) the required non-zero fields around the node display sufficient polarization isotropy required to sufficiently cover the fluorescence dipole orientations in the focus neighborhood. Still, the quality of the zero itself is more crucial than isotropy of the non-zero neighborhood. CH-STED immunity to radial aberrations, along with its background suppression ability, should provide the needed resilient zero, which may prove relevant in imaging dimmer and deeper structures.

## ACKNOWLEDGEMENTS

Work in the laboratory of H.M. is funded by the European Research Council (ERC) under the European Union’s Horizon 2020 research and innovation programme (grant agreement No 681443) and FLAD Life Science 2020. The grant PPBI-POCI-01-0145-FEDER-022122 supports work of M.S. at the i3S Advanced Light Microscopy scientific platform (ALM). A.J.P. is funded by project Norte-01-0145-FEDER-000029, supported by Norte Portugal Regional Operational Programme (NORTE 2020), under the PORTUGAL 2020 Partnership Agreement, through the European Regional Development Fund (FEDER).

## Methods

### Microscope setup

An Abberior Instruments ‘Expert Line’ gated-STED was used coupled to a Nikon Ti microscope. An oil-immersion 60x 1.4NA Plan-Apo objective (Nikon, Lambda Series) and pinhole size of 0.8 Airy units was used in all acquisitions. The system features 40 MHz modulated excitation (405, 488, 560 and 640nm) and depletion (775nm) lasers. The depletion beam (approx. 1ns-long) is modulated by a phase-only SLM (Abberior ‘easy3D’) which allows arbitrary imprinting of phase masks, as well as the incoherent superposition of two arbitrary depletion beams (not used in this study). A 4f-system images the STED beam at the back focal plane of the objective, where a slight overfilling balances doughnut energy with geometrical confinement (NA). To a higher or lesser extent, the effective NA defined by the beam width smoothens the beam-shape response to ρ variation. Because absolute power levels were not required for this study, all STED laser powers were measured integrating the whole beam before hitting the objective. The microscope’s detectors are avalanche photodiode detectors (APDs) which were used to gate the detection (window from0-800ps to 8μs). To prevent saturation, settings (laser power levels, dwell time) were always chosen so that the maximum photo-detection count-rate was below 10MHz.

Abberior’s Imspector software, which is used to control the microscope settings and the whole acquisition process, allows the SLM to be controlled externally via Matlab or Python. A Python script was used to imprint the bi-vortex for CH-STED imaging. The pattern is actually imprinted on top of a factory-set flat-field correction phase map and a grating structure used to diffract the beam off the zero-order diffraction caused by the SLM’s pixelation. The grating period was tuned every couple of hours to warrant co-alignment of the excitation and STED beams. In some images (Fig. 5f, 5i (right panel) and Supplementary Fig. 1), the SLM pattern was switched during scanning (slow scan axis always vertical).

### Beads analysis

xzy-scans of gold nano-beads were performed by detecting the 775nm scattered signal with a PMT positioned before the confocal pinhole with a 20nm pixel size in all dimensions. Depletion beam profiles and concavity determination (Fig. 2a) were done in ImageJ by intensity profiling a 3μm-long 60nm-wide line. Concavity was determined in Matlab by doing a parabolic fit to the eleven datapoints around the profile dip (200nm window). The portion of the SLM imaged on the effective back focal plane of the objective, which must be known for defining the pupil unit circle, is estimated at 108 SLM pixels by inspection of the data in the inset of Fig. 2b.

To characterize the microscope PSF in 2D-STED and CH-STED, 10 fluorescent nano-beads were xz-scanned for each condition (STED power or SLM bivortex radius variation), amounting to a total number of 200 beads. Acquisition pixel size was set to 20nm both dimensions. No acquisitions were excluded. For each bead image, a rough PSF center position was set to allow definition of a relevant ROI matrix (2.8×0.6μm) around the bead image, which was then exported to Matlab for quantification. A Gaussian fit to the x-projected matrix was used to determine a ‘focal plane’ position. The pixel line corresponding to the calculated focal plane was summed to the adjacent lines (defining a 60nm-wide axial averaging) for *FWHM* and *Amplitude* (Figs.4c-e and Fig.4f, respectively) determination by gaussian fitting. The goodness of fit was evaluated using the coefficient of determination *R*^2^, with results higher than 0.90. The same procedure (Gaussian fit after 3-line averaging) was used to determine the out-of-focus (D_z_) width, at a Rayleigh range distance of 260nm (13 pixels) from the focal plane. Background estimation (required to generate Fig. 4f) was done by averaging the full matrix line (600nm-long) at a distance *λ_STED_* from the focal plane. Results are presented as (mean)±(standard deviation) of 10 beads for each condition (Figs. 4c-d).

### Immunofluorescence

Indian muntjac (IM) fibroblasts, immortalized with human hTERT (pBabe puro hTERT, *kind gift from Jerry W Shay*) were grown in Minimum Essential Media (MEM) (Gibco, Life Technologies), supplemented with 10% FBS (Gibco, Life Technologies), 2mM L-Glutamine (Alfagene) at 37 °C in humidified conditions with 5% CO2. IM fibroblast cells were seeded on fibronectin-coated coverslips 2 days before the experiment. Cells were incubated with 20 μM MG-132 (Calbiochem) for 50 minutes before fixation, in order to enrich the mitotic population. After fixation with 4 % Paraformaldehyde and 0.25 % Glutaraldehyde (Electron Microscopy Sciences), the cells were quenched with a 0.1% solution of Sodium Borohydrate in PBS and extracted using PBS-0,5% TritonX (Sigma-Aldrich). The coverslips were incubated with the primary antibodies [anti-human centromere antiserum 1:150 (Fitzgerald); anti-tyrosinated tubulin 1:150 (MCA77G, AbD Serotec)] in blocking solution for 1h at room temperature. After washing with PBS-0.1%Tween, coverslips were incubated with the secondary antibodies (Abberior STAR 580 and STAR RED 1:100) for STED microscopy. DAPI was then added for 5 minutes in PBS-0.1% Tween 1:50,000 (4’6’-Diamidino-2-phenylindole, Sigma Aldrich). Coverslips were washed in PBS and sealed on glass slides using mounting medium (20 nM Tris pH 8, 0.5 N-propyl gallate, 90% glycerol).

Human U2OS were cultured in DMEM (Gibco) supplemented with 10% fetal bovine serum (FBS, Gibco), at 37° C onto 22×22 mm coverslips. Fixation was performed using a solution of 4% (v/v) paraformaldehyde (Electron microscopy sciences) and 0.25% (v/v) glutaraldehyde (Electron microscopy sciences) for 10 minutes. Autofluorescence was quenched using freshly prepared 0.1% (w/v) sodium borohydride (Sigma) for 10 minutes followed by cell permeabilization with phosphate buffer saline (PBS) supplemented with 1% Triton-X100 (Sigma-Aldrich) for 10 minutes. Immunolabeling was performed using rat anti-tyrosinated tubulin (1:200, clone YL1/2, MCA77G, AbD Serotec) and human anti-centromere antibody (1:200, kindly provided by Bill Earnshaw) diluted in PBS supplemented with 20% FBS and fluorescently labelled antibodies anti-rat STAR RED (1:200, Abberior) and anti-human STAR 580 (1:200, Abberior) diluted in PBS supplemented with 20% of FBS. DNA was counterstained with 4’,6-Diamidino-2-phenylindole (DAPI, 1:50 000, Sigma Aldrich). Coverslips were mounted in glass slides using home-made mounting medium (20 nM Tris pH 8, 0.5 N-propyl gallate, 90% glycerol).

Hippocampal neuron cultures from Wistar rats were performed as described (Kaech and Banker, 2006). Briefly, hippocampi from embryonic day 18 pups were digested for 15 min in 0.06% porcine trypsin solution (Sigma, T4799), triturated, and plated at 12,500 cells/well in 24 well-plates containing glass coverslips pre-coated with poly-L-lysine (50 μg/mL; Sigma, P2636). Neurons were cultured in Neurobasal medium (Invitrogen) supplemented with B27 (Gibco), 1% penicillin/streptomycin (Gibco) and L-Glutamine (2 mM; Gibco). At DIV8, cells were washed with PBS, fixed in 4% Paraformaldehyde in PBS (pH 7.4) for 20 min at room temperature, permeabilized with 0.1% Triton X-100 for 5 min, quenched with 200 mM ammonium chloride for 5 min, and blocked in 5% FBS for 1 hour. Primary mouse anti-βII-spectrin (1:200 in blocking buffer; BD Biosciences, cat#612562) was incubated overnight at 4°C and secondary anti-mouse STAR635P (1:200 in blocking buffer; Abberior, cat#2-0002-007-5) was subsequently incubated for 1 hour at room temperature. Incubation with phalloidin STAR635 (300 nM in PBS; Abberior, cat#2-0205-002-5) was performed for 1 hour at room temperature. Coverslips were mounted in 80% glycerol.

HeLa cells were grown in Dulbecco’s Modified Eagle Medium (DMEM) (Gibco, Life Technologies), supplemented with 10% FBS (Gibco, Life Technologies), at 37 °C in humidified conditions with 5% CO2. HeLa cells were seeded on coverslips one day before the experiment. After the fixation with 4 % Paraformaldehyde in PBS pH 7.4 the cells were extracted using PBS-0,3% TritonX (Sigma-Aldrich). After short washes in PBS with 0.1% TritonX and blocking with 10% FBS in PBS with 0.1% Triton X, all primary antibodies were incubated at 4°C overnight (NUPS Abcam 24609 – Mouse 1:100; TPR NB100-2867 – Rabbit 1:100). The cells were then washed with PBS containing 0.1% TritonX and incubated with the respective secondary antibodies for 1h at room temperature (Abberior anti-rabbit STAR580 1:100; anti-mouse STAR 635P 1:100) for STED microscopy. Coverslips were washed in PBS with 0.1% TritonX and sealed on glass slides using mounting medium (20nM Tris pH 8, 0.5 N-propyl gallate, 90% glycerol).

### Imaging sequence and display

Whenever more than one acquisition is shown for the same object, CH-STED was always acquired last (Fig. 3b, Figs. 5b,c,d,e,g,h,i(left)). Whenever a gray-scale bar or color bar is shown, it applies to all images in the associated set. In those image pairs which share the LUT but for which an intensity scale is not shown, a ‘*same LUT*’ label was used explicitly connecting the images. Images acquired by live-shifting the STED mode are a single image, which therefore share the LUT. Chromo-projections (Figs.3a,b and Fig.5d) were created using ‘Temporal Color Code’ in Fiji with the ‘Spectrum’ LUT. Intensity colorbars used in blue-grey-red images use the ‘Phase’ LUT. The independent top-half and bottom-half acquisitions in Fig. 3a were aligned manually by cropping the (common) region in the top half, amounting to a downward shift of 330nm.

## Supplementary Text

### Radial aberrations

Following the seminal work of Richards and Wolf (Proc. R. Soc. Lond. A 1959 **253**, 358-379), we express the electric field components near the focus of an aplanatic lens system as:

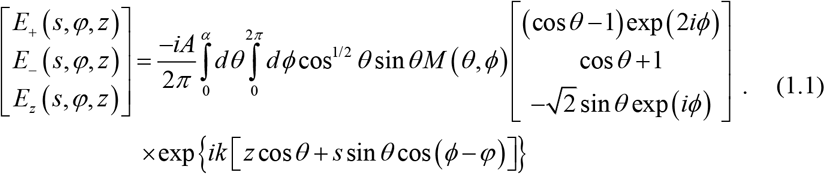

Here we have taken the incident wave to be a right-hand circularly polarized monochromatic plane wave while *M* (*θ, ϕ*) is the phase mask used to generate the STED beam, assumed to be placed at the entrance pupil characterized by the spherical polar coordinates (*θ,ϕ*). The upper limit on the polar angle integration is related to the numerical aperture of the lens system by *n* sin(*α*) = *NA*, while the coordinates in image space are **r** = (*s, φ, z*) centered on the position of the paraxial focus.

The bi-vortex is a member of a family of phase masks that can be called ‘radial vortices’, characterized by a phase *g*(*θ*) + *nϕ*. Here *g*(*θ*) is an arbitrary function that depends only on the polar angle *θ*, and *n* is a non-zero integer defining the vortex order or charge. Within this family one can envision a set of infinitely thin annular rings, each at a given polar angle, modulated by a nth order vortex. In this case the integrals over the azimuthal angle can be written as

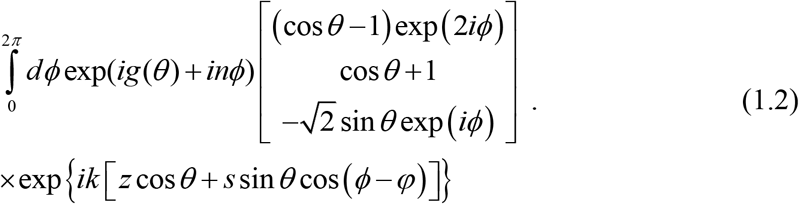

Using the Bessel function identity (Goodman, 2005),

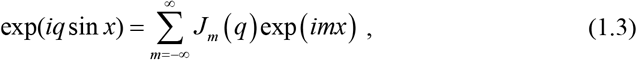

one can readily show that the azimuthal integrals in Eq. (1.2) reduce to

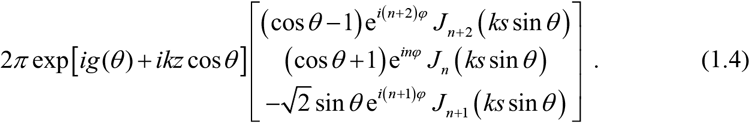

For positive *n*, i.e. when the helicity of the vortex mask is equal to that of the incident circular polarization, all of the Bessel functions have a positive order and vanish on-axis where *s* = 0. Contrary to non-zero disturbances, which may vanish by coherent addition, superimposed zero disturbances will yield an intensity-null irrespective of the nature (coherent or incoherent) of the addition, meaning that the bi-vortex intensity profile will always vanish on-axis irrespective of the type of radial perturbation.

Examples of radial perturbations are:

i. an imprecision in setting c=1 for the bi-vortex cπ-step function,
ii. an imprecision in setting the desired bi-vortex radius ρ,
iii. a spherical aberration term (r^4^),
iv. a paraxial (r^2^) or non-paraxial defocus term on the sample side,
v. a defocus term in imaging the phase mask onto the objective back-aperture.

Of course, spherical aberrations or any radial perturbation may alter the off-axis intensity profile, which can affect the STED depletion in areas not highly saturated.

### Focal plane bi-vortex STED beam profile

To arrive at a simple expression characterizing the focal plane bi-vortex STED beam profile we employ a simplified model, assuming coherent monochromatic plane waves incident on the phase mask placed at the entrance pupil of the lens system. Treating the lens system as a simple thin lens in the paraxial approximation, the cos^1/2^ *θ* ≃ and the incident polarization will be maintained through the image space to a good approximation. Then the electric field amplitude of the STED beam in the focal plane can be estimated within the Fresnel approximation by (Born and Wolf, 2013),

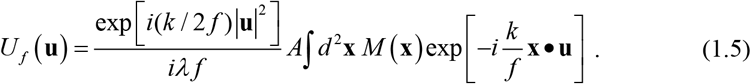

Here, **x** denotes the coordinates in the mask plane, while **u** are those in the focal plane, *f* is the effective focal length of the objective, *A* is the amplitude of the incident plane wave, *λ* it wavelength and *M* (**x**) the bi-vortex mask function:

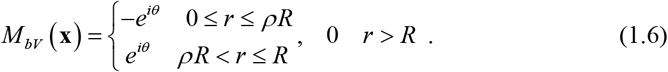

The radius of the circular entrance pupil is *R* while we have used (*r, θ*) to denote the polar coordinates in the mask plane and will take (*s, ϕ*) to be the corresponding polar variables in the focal plane. Within the Fresnel approximation, the electric field distribution in the focal plane is determined by the Fourier transform of the mask function. The Fourier transform of a single vortex function of radius *R* can be shown to be

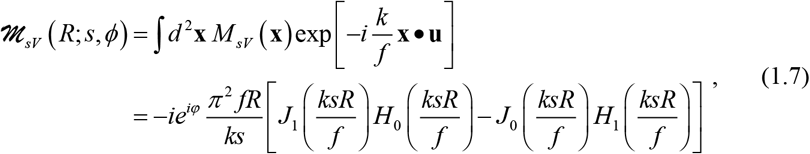

where *J_n_* and *H_n_* are respectively the nth order Bessel and Struve functions (Abramowitz and Stegun, 1965). Here *M_sV_* is equal to *M_bV_* of Eq. (1.6) in the limit that *ρ* → 0 or equivalently *ρ* → 1. Consequently, for the coherent superposition bi-vortex one has

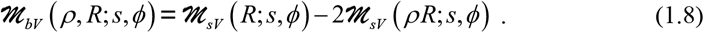

Expanding the resulting expression in a Taylor series about *s* = 0, one can readily show that to leading order in *s*, the focal plane intensity of the bi-vortex near the optical axis plane varies as

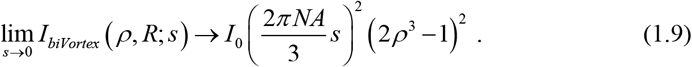

Here we have used the paraxial approximation for the numerical aperture, *NA* ≈ *λkR* / (*2πf*), while *I*_0_ represents the on axis focal plane intensity created by a circular pupil (without a phase mask) of radius *R*. At this expansion level, which we call the parabolic approximation, the only difference relative to the single vortex (2D-STED) mask is the factor of (2*ρ*^3^ − 1)^2^. However, it should be noted that the on-axis curvature of the bi-vortex focal plane intensity pattern becomes increasingly smaller as *ρ* approaches 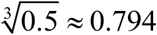 as can readily be seen in Figure A.1.

**Figure A.1:**
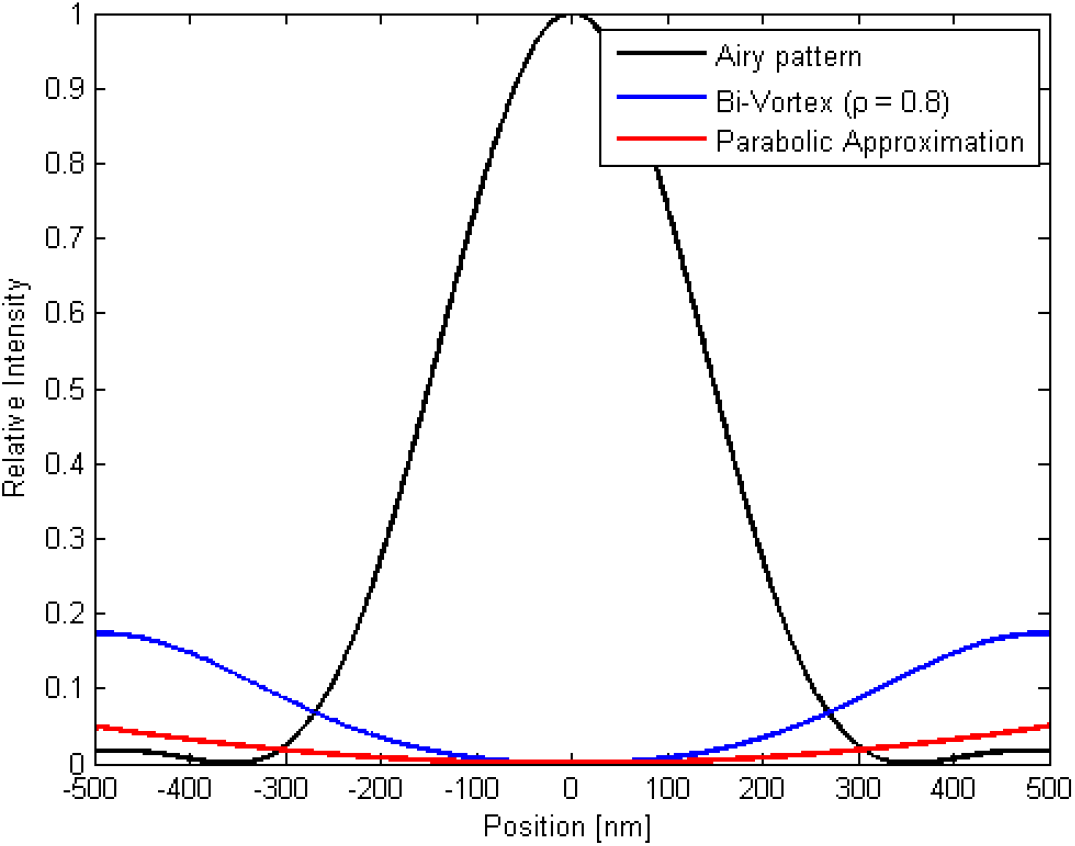
Intensity variation of the bi-vortex focal plane profile near the point at which the on-axis curvature vanishes, together with the leading term in the Taylor expansion (parabolic approximation).

Because of this vanishing curvature, the parabolic approximation to estimate the full width half maximum (FWHM) of the fluorescence profile can lead to significant errors. To arrive at the theoretical curve shown in **Fig. 4d** the following procedure was followed. First the nominal FWHM of the focal plane fluorescence was determined through a nonlinear least squares fit of the experimental profile a Gaussian function, providing a FWHM of 294nm with a 95% confidence interval of [289 300]. This information was used to constrain the fit of the 2D-STED focal plane FWHMs as a function of the STED beam power shown in **Fig. 4c**. We assumed that the convolution with the finite fluorescent beam diameter could be accounted for by adding a constant width in quadrature with the conventional 2D-STED FWHM power dependence:

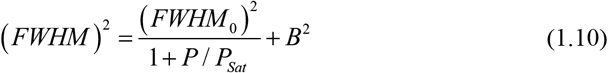

Using a weighted nonlinear least squares fitting routine with the nominal FWHM constrained to lie within the above 95% confidence interval of the focal plane fluorescence profile, we obtained the following parameters *FWHM*_0_ = 300 *nm, P_sat_* = 5 *mW, B* = 32*nm* with an adjusted R-squared value of 0.995. To obtain the FWHM profile for the CH STED we assumed a simple rate equation model for the excitation and de-excitations of the excited fluorescing molecular state. In the focal plane this leads to an emitted fluorescence profile of the form

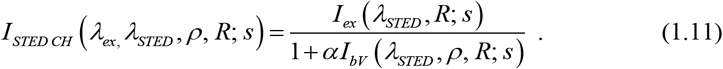

Here *I_ex_* (*s*) represents the excitation profile (Airy function), *I_bV_* (*s*) the bi-vortex intensity profile corresponding to

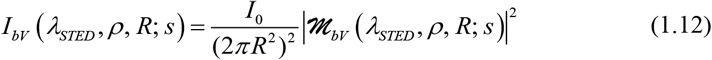

The parameter *α* was set by requiring that the FWHM of the profile given by Eq. (1.11) with *ρ* = 1 correspond to that measured using the 2D-STED configuration with an incident power of 60mW, yielding *α* = 59. Numerically determining the resulting FWHM from Eq. (1.11) as *ρ* is varied and including the bead convolution contribution B added in quadrature, leads to the curve shown in Figure A.2 reproduced as the theoretical curve of **Fig. 4d**. For comparison, the prediction of the conventional expression for FWHM reduction based on the on-axis curvature (the parabolic approximation of Eq. (1.9) is also included.

**Figure A.2:**
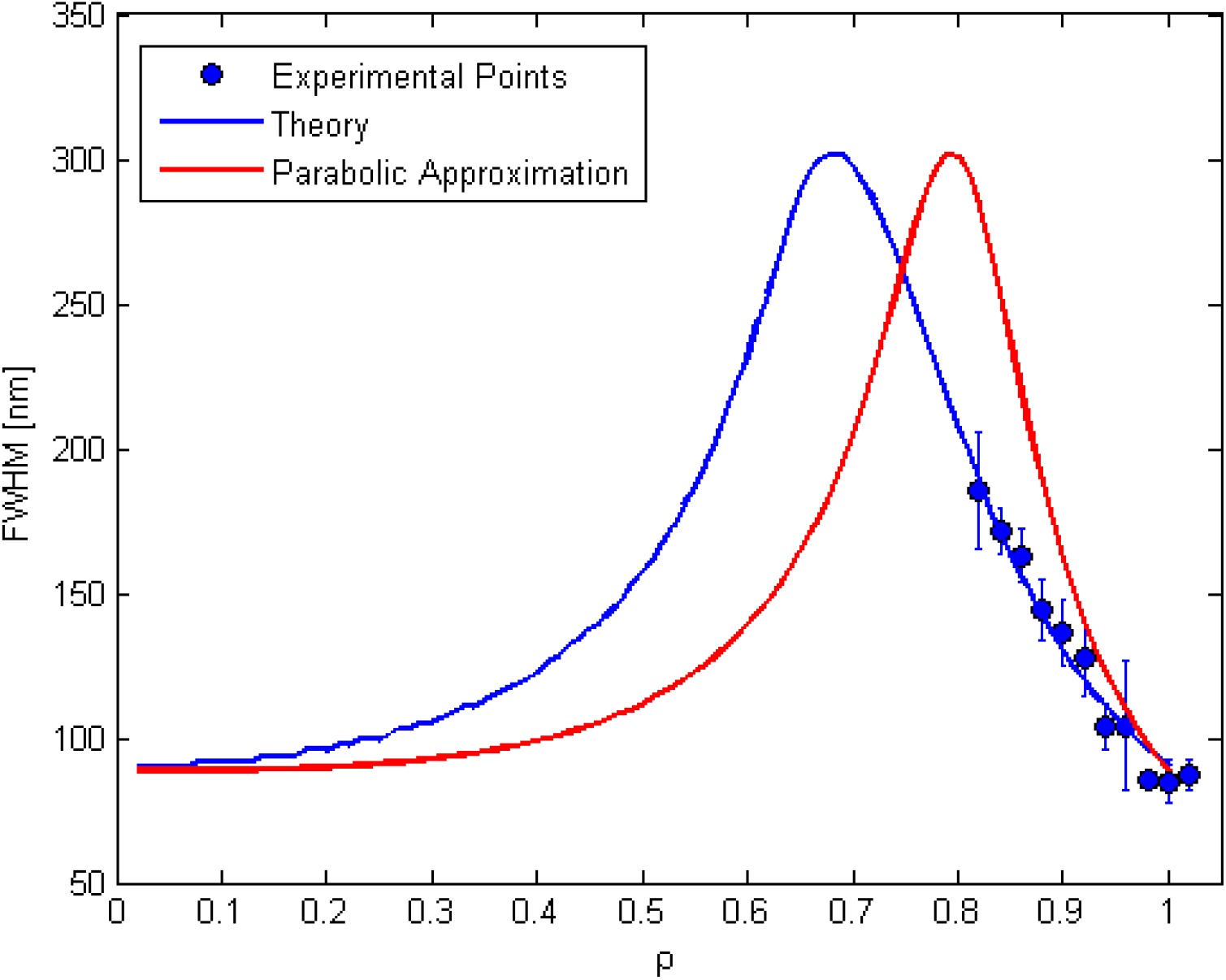
Predicted variation of the focal plane fluorescence spot size as the STED CH parameter *ρ* is varied. The parabolic approximation deviates significantly from the experimental values for values of *ρ* below 0.9.

## Supplementary Figure Legends

**Fig. S1**. Switching between different STED modes while scanning a field of Indian muntjac interphase cells. Apart from the STED beam shape, all acquisition settings are kept constant. The first mode (second-order vortex) is an archetypal dip enlargement strategy that provides signal rescue but, just as when decreasing STED power, it rescues defocused signal also. z-STED displays notorious sectioning, at the expense of lateral resolution and SNR.

## Acronym list

CH: coherent-hybrid
LUT: look-up table
PSF: point-spread function
SLM: spatial light modulator
SNR: signal-to-noise ratio
STED: stimulated emission depletion

